# THE MATRIX IS EVERYWHERE: CACO_3_ BIOMINERALIZATION BY THE *BACILLUS LICHENIFORMIS* PLANKTONIC CELLS

**DOI:** 10.1101/2020.10.22.351619

**Authors:** Lyubov A. Ivanova, Darya A. Golovkina, Elena V. Zhurishkina, Yuri P. Garmay, Alexander Ye. Baranchikov, Natalia V. Tsvigun, Yana A. Zabrodskaya, Alexey D. Yapryntsev, Andrey N. Gorshkov, Kirill I. Lebedev, Aram A. Shaldzhyan, Gennady P. Kopitsa, Vladimir V. Egorov, Anna A. Kulminskaya

## Abstract

To date, the mechanisms of CaCO_3_ nucleus formation and crystal growth induced by bacterial cells still remain debatable. Here, an insight on the role of planktonic cells of *Bacillus licheniformis* DSMZ 8782 in the biomineralization is presented. We showed that during 14-days bacterial growth in a liquid urea/Ca^2+^-containing medium the transformation of CaCO_3_ polymorphs followed the classical pathway “ACC-vaterite-calcite/aragonite”. By microscopic techniques, we detected the formation of extracellular matrix (ECM) around the cells at the stage of exponential growth and appearance of electron-dense inclusions at 24 h after the inoculation. The cells formed filaments and created a network, the nodes of which served as sites for further crystal growth. The ECM formation accompanied with the expression of proteins required for biofilm formation, the aldehyde/alcohol dehydrogenase, stress-associated Clp family proteins, and a porin family protein (ompA ortholog) associated with bacterial extracellular vesicles. We demonstrated that urea and CaCl_2_ acted as denaturing agents causing matrix formation in addition to their traditional role as a source of carbonate and Ca^2+^ ions. We showed that CaCO_3_ nucleation occured inside *B. licheniformis* cells and further crystal growth and polymorphic transformations took place in the extracellular matrix without attaching to the cell surface. The spatial arrangement of the cells was important for the active crystal growth and dependent on environmental factors. The extracellular matrix played a double role being formed as a stress response and providing a favorable microenvironment for biomineralization (a high concentration of ions necessary for CaCO_3_ crystal aggregation, fixation and stabilization).

## 1. INTRODUCTION

Biomineralization is the phenomenon of mineral formation by living organisms within their metabolic reactions with the environment. Calcium carbonate is one of the most widespread material on the Earth, being the main constituent of limestone, marble, chalk, travertine, dolomite. In the world’s oceans its content reaches 10%. It is an essential component in biological systems such as shells of marine organisms, pearls, and eggshells. Besides that, numerous microorganisms are involved in the calcium carbonate mineralization. The process of microbially induced calcium carbonate precipitation (MICP) has received a lot of attention from researchers working in various fields. Thus, one of the most intensively studied subjects is the development of technologies that use microorganisms able to induce the precipitation of calcium carbonate to restore concrete structures. In the most cases, the microbiological method has been developing for the restoration of historical stone buildings (1, 2). The main advantage is the imitation of the natural mineral formation that guarantees good compatibility of the newly formed CaCO_3_ mineral and the building material (2, 3).

Three anhydrous polymorphs of crystalline calcium carbonate (aragonite (4), calcite (5) and vaterite (6)) and two hydrated phases (monohydrocalcite and ikaite) (7) are well-described. In biogenic and non-biogenic calcium carbonate minerals, the amorphous calcium carbonate (ACC) phase is also found and has been received increasing attention by researchers (8). The most thermodynamically stable CaCO_3_ polymorph is calcite, the least stable non-crystalline polymorph is ACC (9, 10). The latter often serves as a precursor in the formation of more stable crystalline polymorphs (11). Recent works report various applications of CaCO_3_ polymorphs including ACC to create new biomaterials (12). As a consequence, morphological control of crystal growth is necessary during the development of new products with unique characteristics (13).

To date, the mechanism of crystal formation by bacterial cells still remains debatable. In a recent study (14), Keren-Paz and Kolodkin-Gal suggested a possible scheme of biogenic CaCO_3_ crystallization pathway. The authors offered to divide the process into 4 stages: a) formation of a saturated CaCO_3_ micro-environments; b) nucleus formation; c) crystal nucleus transfer out of the cell; and d) crystal growth on or near the cell surface. The authors supposed that at the first stage, Ca^2+^ is accumulated in a cell as a result of ATP-dependent calcium transporters action or as a product of carbonic anhydrases function within caboxysomes. During the second stage the initial nuclei are forming and accumulating to be further transferred and to evolve at the third and fourth stages. Very often, nucleation is thought to occur on microbial membranes. It is expected that viable cells cannot produce crystals larger than their size; accordingly, at the third phase the formed nuclei are transferred to the surface of the bacterial cell using active transport or lysis. At the last stage, crystal growth occurs through the aggregation of nuclei on charged cell walls. The rate, at which a crystal grows, can be affected by the shape of the crystal and its morphology, the environment, and external conditions. Despite the rather logical proposed scheme of biogenic crystal formation, a large number of questions arise at all stages. For example, nucleus formation and initial crystal growth are the most questionable issues. To date, there are a lot of contradictory results obtained regarding the influence of the microbial cell and its closest environment on mineral formation.

Several research groups reported the intracellular carbonate ion formation within metabolic reactions of the cell together with free calcium ions attraction out of the cell followed by the formation of insoluble CaCO_3_ crystals at the negatively charged cell wall surface (15). In this case, the crystal morphology depends on many factors: the bacterial cell morphology, environmental components, as well as physical factors (temperature, pH, and aeration) (16). This was confirmed by Ghosh et al. who demonstrated appearance of nanoscale calcium carbonate crystals at the surface of *Sporosarcina pasteurii* (17) supporting the hypothesis of the critical role of a negatively charged surface in the formation of crystals. In contrast, W. Zhang et al. claimed that the mechanism of crystal formation proceeds according to a different scheme (18). In addition, Bundeleva et al. (19) reported studies on anoxygenic *Rhodovulum* sp., in which no crystals were observed on the surface and near the cell. The authors hypothesized about the mechanisms of cell protection against mineral deposition, and they proposed the idea of crystal deposition at some distance from the cell surface. Zhang C. et al. (20) assumed that it is a spherical core formed at the initial stage followed by the formation of a crystal structure with various shapes depending on the selected conditions. The amorphous phase (ACC) also plays an important role, being one of the main components of the spherical core (12). Thus, the question of the bacterial cells role remains unexplicit and still requires a more detailed study of the exact mechanism of calcium carbonate bioprecipitation.

In previously performed screening for the bacterial strains able to MCIP we found that one of the most potential cultures was the strain *Bacillus licheniformis* DSMZ 8782 (results will be published elsewhere). In the present study we used several microscopic and biochemical techniques to demonstrate the behavior of planktonic bacterial cells during the growth in urea-containing medium and their effect on initiation of CaCO_3_ crystallization and crystal evolution.

## 2. RESULTS

### 2.1. Particle analysis produced by *B. licheniformis* DSMZ 8782 during 14 days

Samples with CaCO_3_ precipitates obtained during the growth of the bacterium in the liquid medium B4-UCa for 14 days were analyzed by FTIR, XRD and SEM methods (Fig. 1). Intense absorption bands of the carbonate ion (ν_1-4_ CO_3_^2-^), OH-groups and water (21) can be easily distinguished on the FTIR spectra of the obtained samples (Fig. 1**A**). Low-intensity bands with maxima at 1800 and 2500 cm^−1^ can be attributed to the composite vibrational frequencies of the carbonate ion or organic impurities (ketones, thiols, or amines) (22). Intense bands ν_2-4_ CO_3_^2-^ (~710, ~870 and ~1400 cm^−1^) correspond to the most abundant calcite phase in the obtained samples. However, the presence of the ν_1_ CO_3_^-2^ band (~ 1080 cm^−1^) indicates the presence of small impurities of aragonite, vaterite, or amorphous calcium carbonate (ACC) in the studied samples (23, 24). The band with a maximum of ~ 750 cm^−1^ has been found only in the spectra of samples collected at points 30 and 48 hours after the inoculation indicating the presence of vaterite (24, 25) in their composition. The amorphous phase is usually characterized by the ratio of the ν_2_ to ν_4_ band intensities for the carbonate ion (20). Accordingly, this ratio increased in the series of samples: at 14 days, 7 days, 48 h, and 30 h after inoculation and reached a maximum for samples containing traces of vaterite phase. Thus, the data of FTIR spectroscopy are in a good agreement with the known model of calcium carbonate polymorphs transformation: ACC → vaterite → calcite, aragonite (25).

**Fig. 1.**
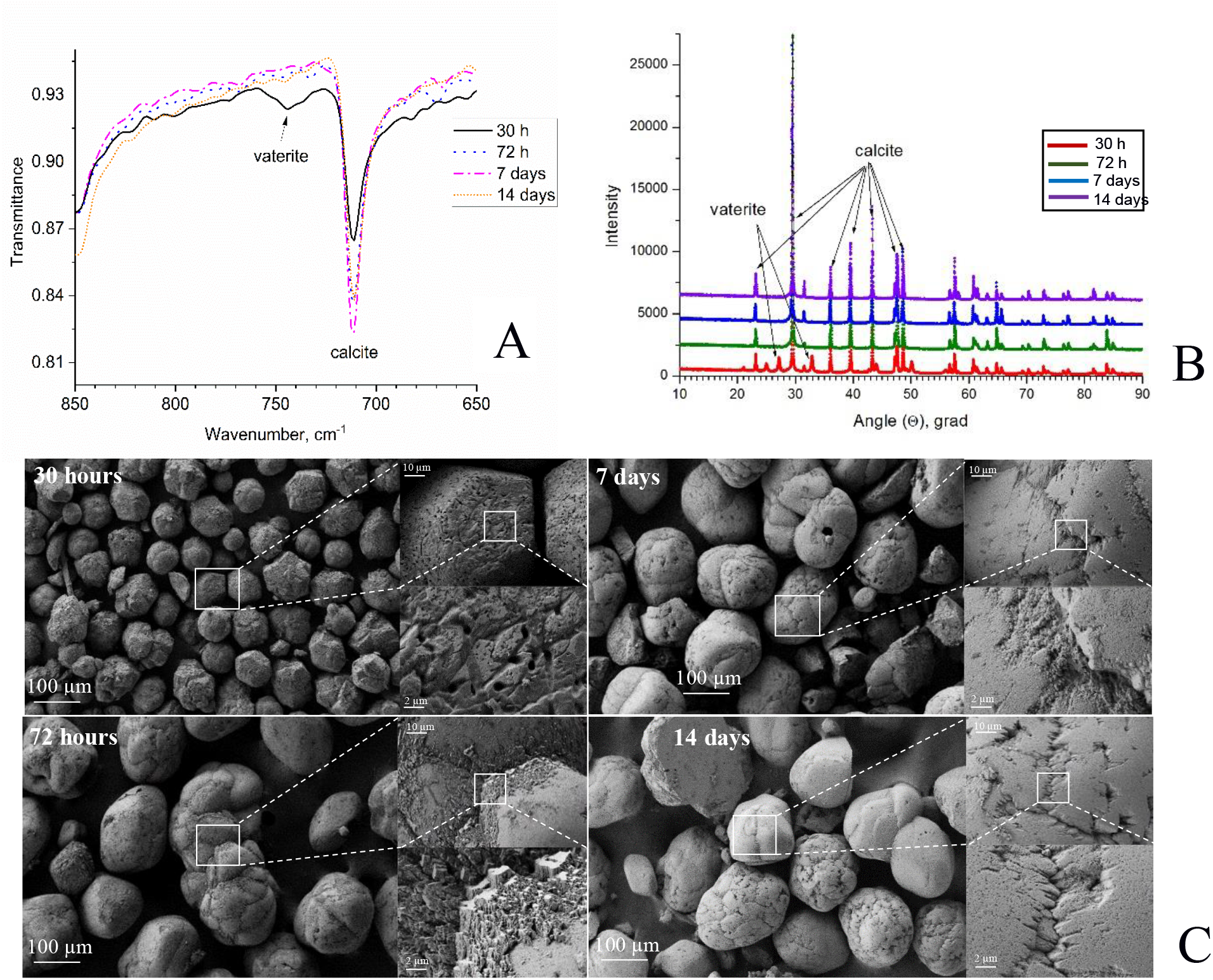
Mineral analysis of CaCO_3_ precipitates induced by the *B. licheniformis* DSMZ 8287 in the liquid medium B4-UCa for 14 days. FTIR spectra of precipitates at different times of growth shows vaterite peak loss after 30 hours: **A**, X-ray diffraction patterns of the samples shows polymorph transition from vaterite to calcite: **B**, variation of SEM topologies, particles growth and loss of amorphous environment by 14^th^ day: **C**.

The X-ray diffraction analysis (Fig. 1**B**) revealed changes in the distribution of calcium carbonate polymorphs during the microorganism growth. Therefore, for precipitated samples washed out from the culture liquid at the point 30 hours after the *B. licheniformis* inoculation, a noticeable amount of vaterite, consisting about 10% of the total content of calcium carbonate crystals, was detected. Further, the vaterite phase gradually transformed into calcite so that in the samples at 72 hours after the inoculation the vaterite peaks were no longer detected. Accumulation of calcite phase was accompanied with ACC disappearance detected by scanning electron microscopy (Fig. 1**C**). At the point of 30 h after the inoculation small spherulites (50-70 μm in size) with bacterial cell traces on their surfaces were located among the amorphous phase as seen in the sample microphotographs (Fig. 1**C**: left top image). After 72 hours of cultivation, the inorganic precipitates increased in size reaching a structure with an average diameter from 300 to 500 μm (Fig. 1**C**: left bottom image). Noteworthy, the smaller precipitates had better crystal structure than their 30-hour precursors. During the following period, the precipitates hardly increased in size, their structures looked crystalline, their amorphous environment disappeared completely (Fig. 1**C**: right).

### 2.2. Analysis of the *B. licheniformis* DSMZ 8782 growth and biomineralization during the first 48 hours

The most intriguing questions in understanding the processes of the calcium carbonate biomineralization is the role of bacteria in the mineral formation. To address this question, we conducted a series of experiments using several different microscopic and biochemical techniques observing the cell behavior during the first 48 hours of the bacterium growth. Another reason to choose this period for further studies was the absence of significant changes in cell behavior and noticeable crystal transformation according to the results described above.

#### 2.2.1. Growth parameters

To study the process of mineral formation induced by *B. licheniformis* DSMZ 8782 in details, pH parameters, biomass accumulation and the specific urease activity were monitored during the bacterial growth for the first two days in the B4-UCa medium (Fig. 2).

**Fig. 2.**
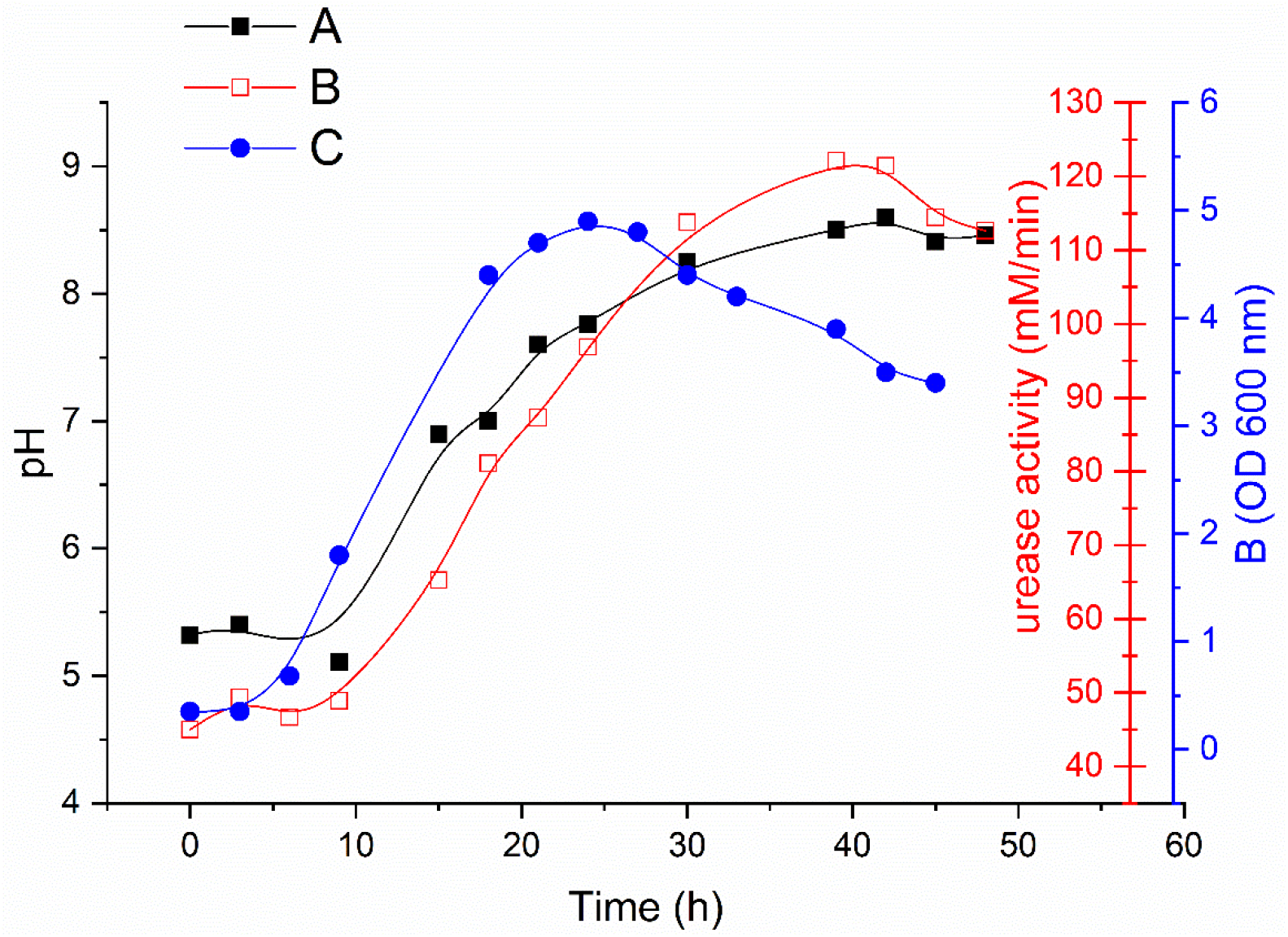
Time-dependences of *B. licheniformis* DSMZ 8782 growth parameters in the B4-UCa medium for the first 48 hours. **A:** pH; **B:** specific urease activity; **C:** biomass accumulation.

After the first 24 hours of the bacterium growth, we observed an increase of the urease activity along with cell biomass accumulation and a subsequent increase in the medium pH. Then, changes in the parameters were stabilized and after about 40 hours the level of the urease activity and cell biomass slightly decreased. We supposed that this was due to the formation of the calcium carbonate precipitates (8).

#### 2.2.3. Light microscopy and AFM

The initiation of CaCO_3_ crystal formation was monitored by the light and atomic-force microscopies (Fig. 3) using samples taken each 3 hours during the bacterial growth in the control (B4-C, urea- and Ca^2+^-free) and the experimental (B4-UCa, with the excess of urea and Ca^2+^) media. At the point 0 h separated cells were seen in the images of the control samples (Fig. 3**A**), followed by their apical collection in 2-4 cell straight chains. By 9 hours of inoculation (Fig. 3**C**), further chain elongation tended to create an untight network. By the same time (9 hours after inoculation), cells in the B4-UCa medium have formed long twisted chains and loops followed by disappearance of clear contacts between pairs of cells and the appearance of amorphous surrounding around bacterium by 15 hours of the growth (Fig. 3**B** and **D**). We supposed this amorphous surrounding to be an extracellular matrix (ECM). At the 24-h point, the images of the samples grown in two media were completely different (Fig. 3**E** and 3**F**): in the B4-C medium separate cells and their stacks were still observed with hardly detectable extracellular matrix in the places of cell accumulation (Fig. 3**E**). The light microscopy and AFM images of the cells in B4-UCa medium demonstrated clearly visible ECM around a cluster of cells with “early” CaCO_3_ crystal inclusions in the network nodes (Fig. 3**F**). By 39 hours of observation, in the images of the control samples separate spores and their accumulations were seen exclusively (Fig. 3**G**), while in the experimental sample the cells have continued to form dense structures around themselves until the height of the “early crystals” reached the maximum to be hardly detected by atomic-force microscopy (Fig. 3**H**). However, the bacterial cell network with numerous crystal precursors were still good visible by light microscopy showing a sharp increase of their number (Fig. 3**H**).

**Fig. 3.**
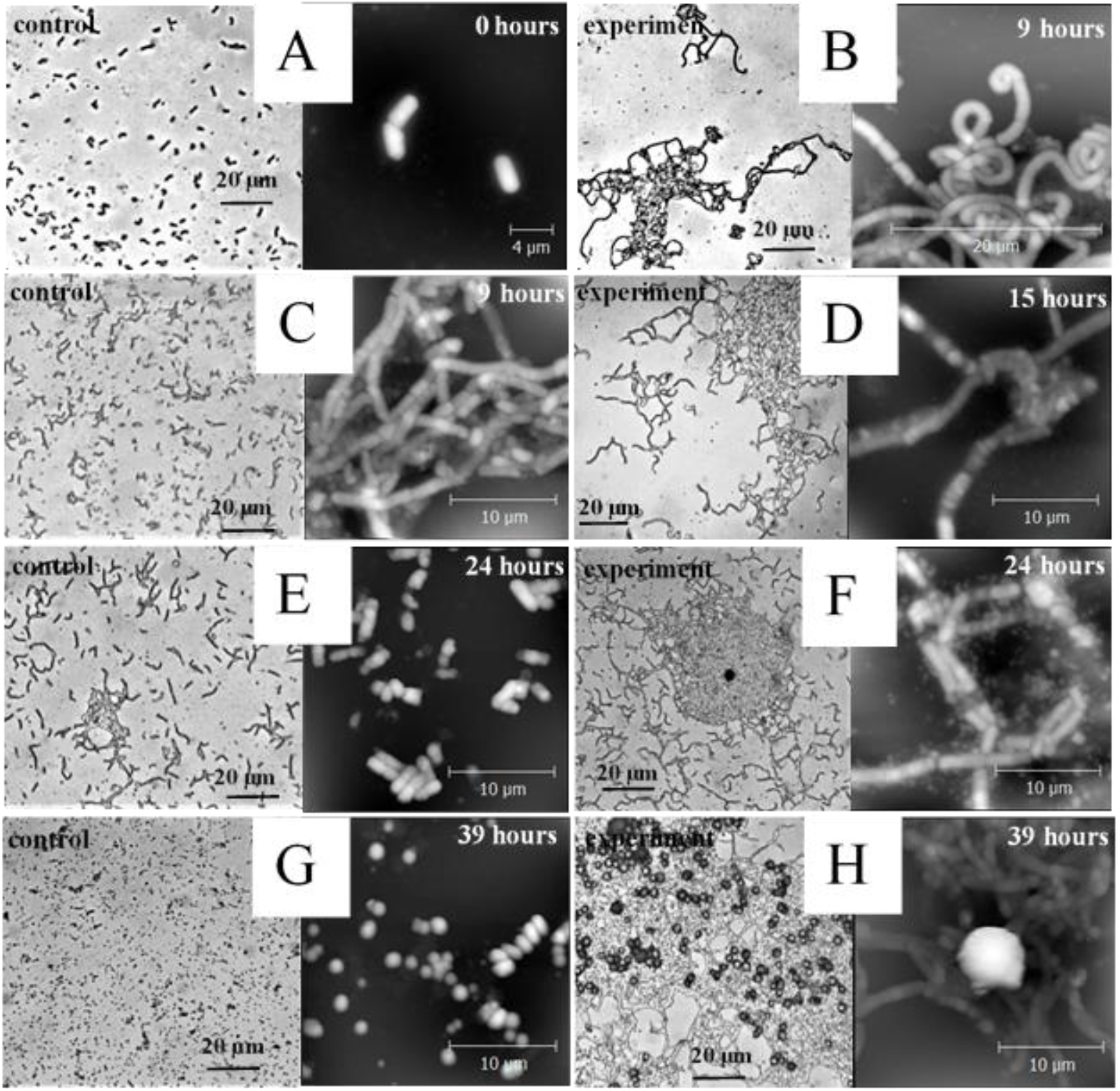
*B. licheniformis* cell morphology evolution during the first 48 hours of the growth. Images of bacterial cells grown in the B4-C (control: **A, C, E, G** pictures) and B4-UCa (experiment: **B, D, F, H**) media were obtained by the light (left) and atomic force (right) microscopies.

The main result of the light microscopy and AFM observations was a clear increase in the amount of extracellular matrix around the cells during the cultivation in the B4-UCa medium, while this phenomenon was not observed for the samples in the urea- and Ca^2+^-free media during the same period. Observations of the cell behavior in both media revealed two trends. First, in an environment with urea and calcium, the life cycle of cells was longer, and the cells formed a strong network, in the nodes of which crystal nucleation occurred followed by the subsequent active crystal growth. In the B4-UCa medium, cells combined into longer chains as compared to the control B4-C medium; their terminal parts were twisted into loops not observed in the control samples. In the control B4-C medium, spores have started to appear 18 hours after the inoculation and their number increased significantly during the studied period. No precipitates were detected in this case. Second, we found the appearance of an extracellular matrix around planktonic cells after 9 hours of the growth in B4-UCa medium. This is typical for the representatives of the *Bacillus* genus including *B. licheniformis,* though it was shown during the growth of microorganisms on a solid medium and was associated with biofilm formation (see for example, (26) or (14)). These observations tended us to find out, which components of the B4-UCa medium make the cells to behave unusually in a liquid medium forcing them to form a specific network, at the nodes of which CaCO_3_ crystal nucleation and growth occurs. To understand what may affect mineral formation in planktonic cells, analysis of the protein content of the extracellular matrix of *B. licheniformis* was required.

### 2.3. Effect of medium components on CaCO_3_ crystallization

To study the influence of the medium composition on the cell behavior and the formation of the extracellular matrix, the following media were used: B4-C, control medium without urea and calcium ions; B4-U contained only urea; B4-Ca contained calcium ions and the experimental medium B4-UCa contained urea and Ca^2+^ (CaCl_2_). Changes in biomass were monitored during the growth of bacteria in all media over a 72-hour period. In B4-U and B4-C media the strain *B. licheniformis* DSMZ 8782 showed similar growth rate reaching their peak after 10 hours of cultivation, after which some decrease was seen (Fig. S1, Supplementary material). One can assume that urea served as an additional nutritional source for the bacterium. Slower growth with a peak at 20 hours was observed in the experimental medium B4-UCa. After 24-hour growth in this medium, intensive crystal formation started to be visible that interfered with density measurement and made it difficult to evaluate biomass accumulation. By the second day of cultivation, amount of *B. licheniformis* cells reached approximately the same values in all media and then their growth was constant.

We selected a point of 24 hours of the bacteria cultivation to make light and atomic force microscopy images for the sample withdrawn from studied media (Fig. 4). Cells in control B4-C medium behaved disassociately gathering into clusters of separable cells (Fig. 4**A**, left). At the 3D-AFM image (Fig. 4**A**, right) the absence of extracellular matrix around bacterial cells were clearly seen. In the presence of urea in B4-U medium (Fig. 4**B**), dense clusters of poorly distinguishable cells were observed which apparently formed a dense extracellular matrix. The excess of calcium ions (B4-Ca medium, Fig. 4**C**) forced the bacterial cells to apically coalesce into twisted chains with coiled strands. Most likely, this cell behavior is associated with the redistribution of ions on the membrane and was also observed in the complete B4-UCa medium (Fig. 4**D**). The AFM image of cells in B4-Ca medium (Fig. 4**C**, right) also demonstrated the accumulation of ECM around bacterial cells, however, in a smaller amount than in the presence of urea (Fig. 4**B**). The light-microscopic image of the sample from the medium B4-UCa (Fig. 4**D**, left) demonstrated a combination of the patterns observed for B4-U and B4-Ca media: the twisted chains of bacterial cells were obviously surrounded by dense extracellular matrix. On the AFM 3D-image of this sample (Fig. 4**D**, right), individual cells were indistinguishable from each other in the extracellular matrix and inclusions of calcium carbonate were clearly observed.

**Fig. 4.**
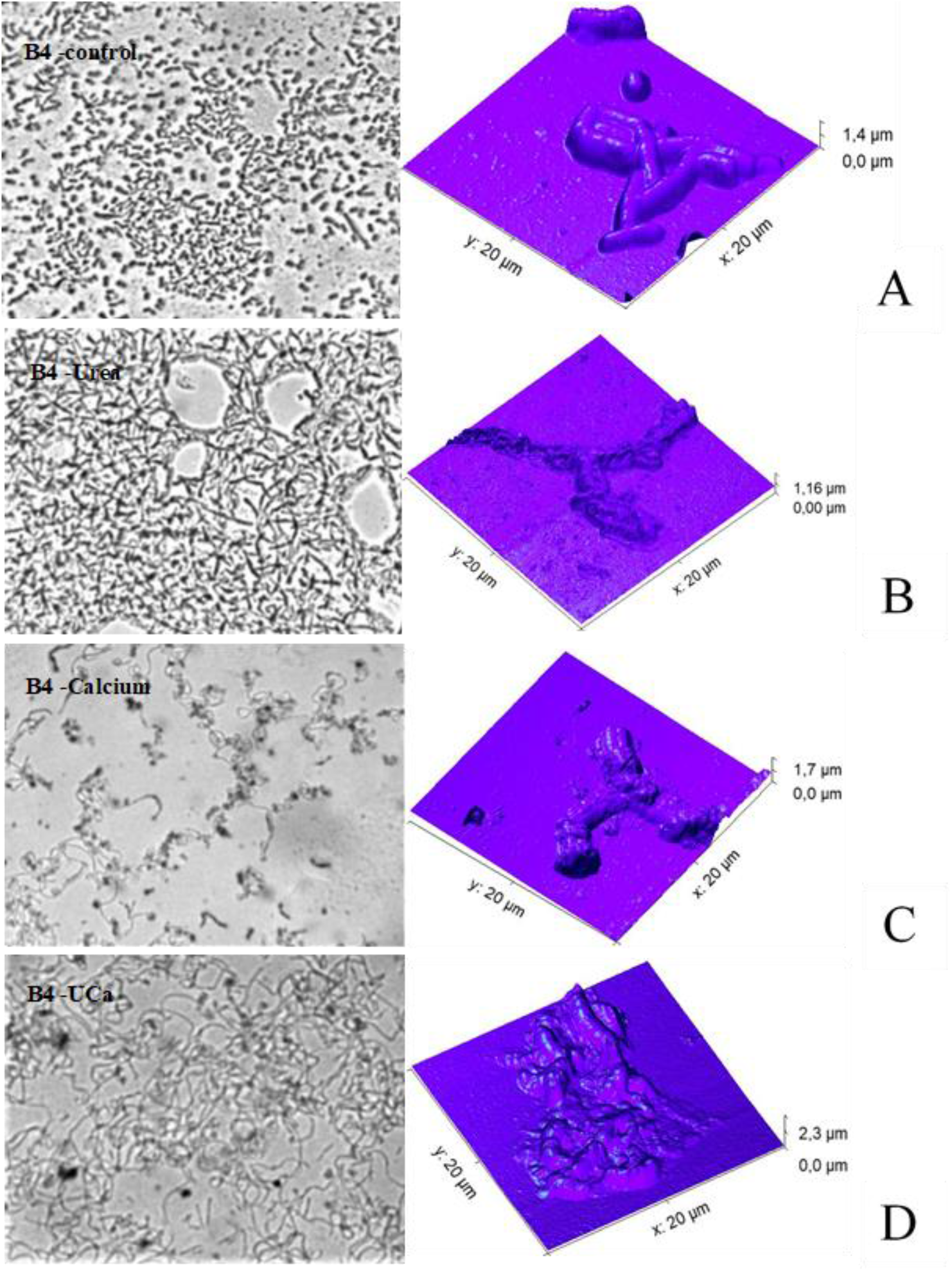
*B. licheniformis* cell morphology at the 24 hours after inoculation. Bacterial cells were grown in four media: the B4-C (control, **A**), B4-U (with urea, **B**), B4-Ca (with Ca^2+^, **C**) and B4-UCa (with urea and calcium, **D**). Samples were observed by the light (left) and atomic force (right) microscopies.

### 2.4. ECM proteome analysis

To analyze differences in proteome composition in ECM of cells grown in the control B4-C (urea- and Ca^2+^-free) and full B4-UCa media, the protein fractions obtained from PBS-washed cells containing extracellular matrix (fractions C, see Materials and Methods, sections 4.4-4.5) were analyzed by PAGE separation (Fig. 5**A**) followed by mass-spectrometric identification (Figures S2, S3, S4, supplemental materials) and densitometric analysis (Fig. 5**B-D**). Analysis of the experimental cell washes revealed a protein zone practically absent in the control (Fig. 5**A**) and accumulated during the bacterium growth. The protein content of another two zones for the experimental cells was more than twice higher than in the control cells, which was most pronounced for the samples corresponding to 15 hours after the inoculation (Fig. 5**A**).

**Fig. 5.**
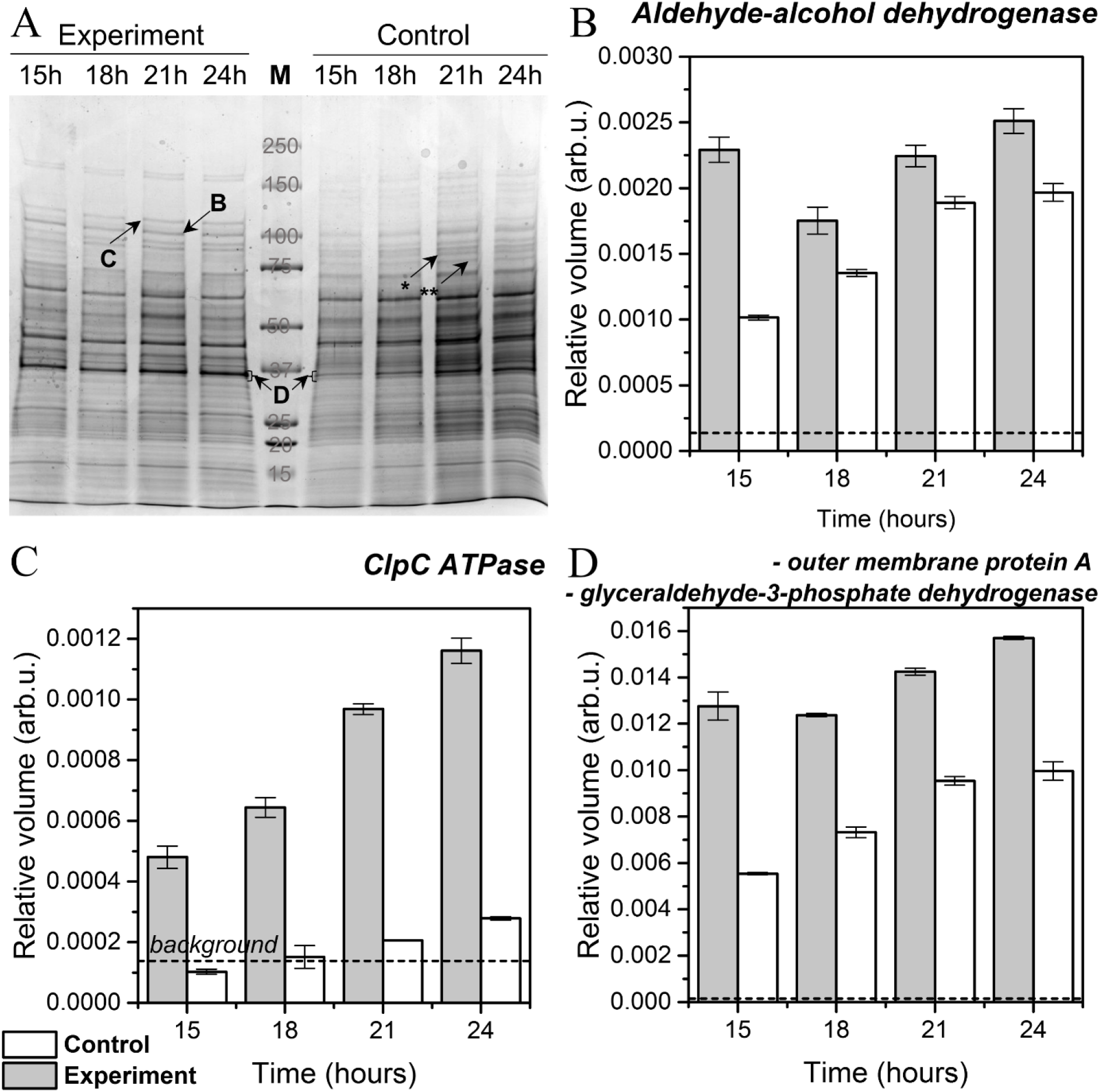
Proteome analysis of the ECM washed with cells during the growth for 24 h. **A:** image of a protein PAGE separation of the fraction C (PBS-washed cells and ECM) produced by the cells in the control and experimental media; **B – D:** densitometric analysis of the zones differed in color intensity (marked with arrows on the panel **A**). The relative volume values were obtained by normalizing the color intensity of a zone to the total color intensity of the corresponding lane. The background value of the relative intensity (indicated by the dashed line) corresponds to the normalized random area of the corresponding track of a similar size.

Homology search with the BLAST software (https://blast.ncbi.nlm.nih.gov/Blast.cgi) revealed about 80% homology between ATPase ClpC, aldehyde-alcohol dehydrogenase from *Enterobacter cloacae* and *B. licheniformis* proteins, and outer membrane protein A from *E. cloacae* mixed with glyceraldehyde-3-phosphate dehydrogenase identified using mass-spectrometric databases. The discrepancy between the amino acid sequences of some of the peptides, for which the identification of proteins was carried out, can be explained by the poor presentation of the amino acid sequences of Gram-positive bacteria, in particular, *B. licheniformis,* in the NCBI and SwissProt databases.

### 2.5. TEM analysis

Electron microscopy of ultrathin slices of *B. licheniformis* cells showed homogeneous, fine-grained with a moderate electron density cytoplasm of the control cells. Rounded vacuoles corresponding to the invaginations of the bacterial membrane were regularly detected (Fig. 6**A**). In experimental cells a higher electron density of the cytoplasm was observed indicating the differences in the chemical composition of the intracellular contents. In the almost all studied experimental samples, rod-shaped, branching dense formations were clearly seen in the whole cell volume and especially along the edge of the inner membrane (Fig. 6**B**, marked by black arrows).

**Fig. 6.**
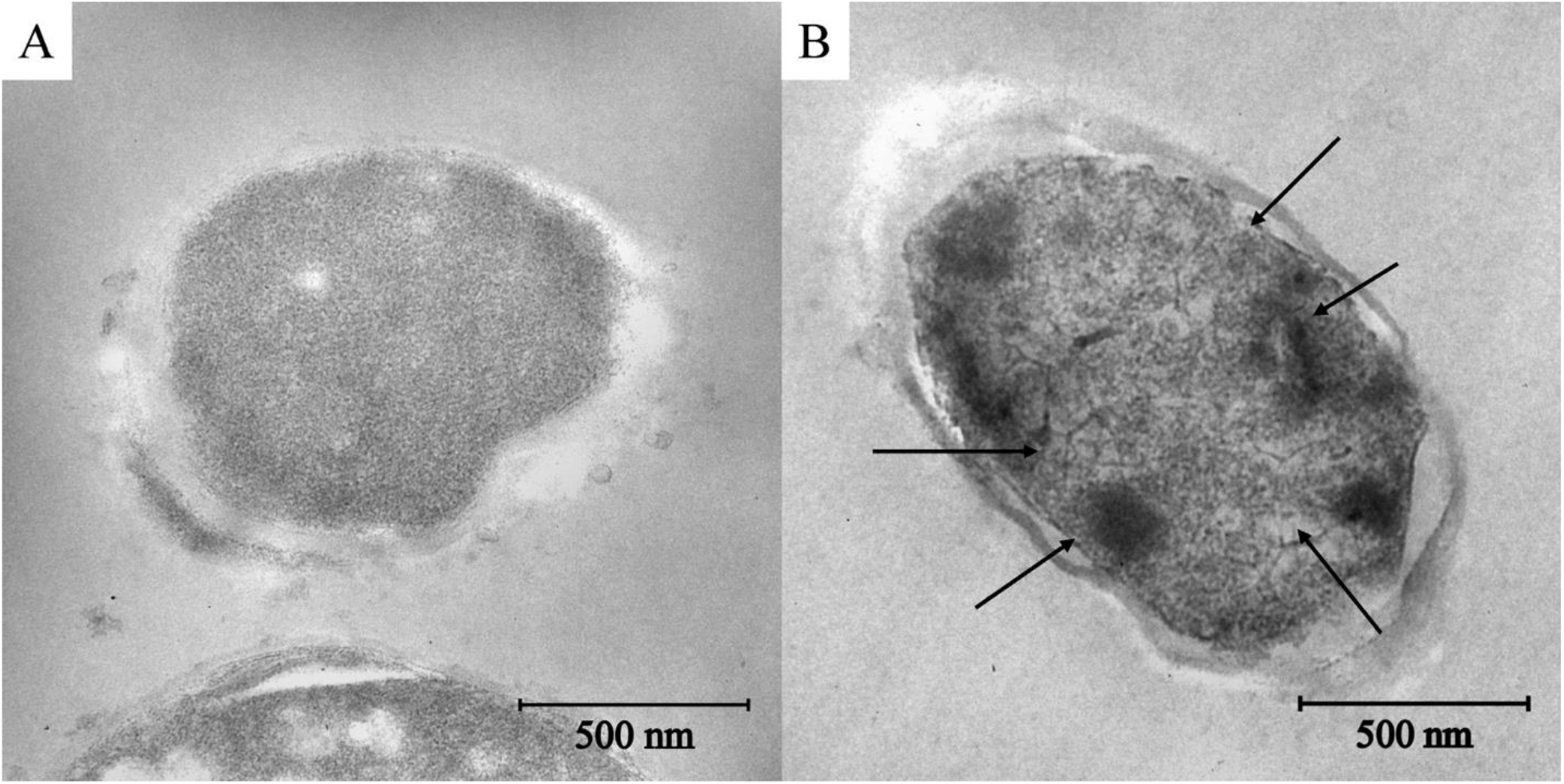
Transmission electron micrographs of unstained cells of *Bacillus licheniformis* at 24 h after the inoculation. **A:** a sample withdrawn from the control medium B4-C; **B:** a sample from the B4-UCa medium. Electron dense inclusions within the cell are indicated by black arrows.

## 3. DISCUSSION

At present a few research groups study intensively the biogenic process of CaCO_3_ mineralization induced by microorganisms, in particular, by representatives of the genus *Bacillus,* as well as the relationship between mineral and biofilm formation (14, 27, 28). However, there are practically no works devoted to the behavior of planktonic cells during the biomineralization induction. Our previous screening for bacteria capable of inducing calcium carbonate precipitation resulted in a choice of the *B. licheniformis* strain DSMZ 8287 as the most promising microorganisms to induce mineral formation (results will be published elsewhere). Our purpose in the present work was to deepen the understanding of the role of planktonic bacterial cells in the biomineralization process. We have carried out detailed studies of the induction of CaCO_3_ precipitation by the studied strain during its 14-days growth in a liquid medium with urea and calcium salts. Under the conditions of our experiments with this strain, the calcium carbonate crystal nucleation, most likely, occurred in the first 24 hours, then the action was transferred to the extracellular space, where the growth of crystals became chemically controlled depending on the physico-chemical parameters of the microenvironment. Our observations revealed that the most important events took place during the first two days, then the changes in the quantity and quality of the formed minerals occurred at a noticeably lower rate and were not so significant. Like it has been reported previously for other bacteria (28, 29) the transformation of CaCO_3_ polymorphs induced by *B. licheniformis* DSMZ8782 followed the classical pathway “ACC-vaterite-calcite/aragonite”. Interestingly, the polymorph transformations resembled a stepwise mechanism of the formation of a substance characterized by several polymorphic modifications, according to the rule of Ostwald steps (30). Although we have not had satisfied data to state this with certainty.

Our observations of the events occurring in the reaction medium with cells in course of crystallization induction for the first 48 hours prompted us to investigate this period in more detail. Analysis of cell images made every 3 hours for the first 48 hours of the bacterial growth in the experimental (with an excess of urea and calcium salts) and in the control (urea/Ca^2+^-free) media led us to the following conclusions. At the stage of exponential growth, ECM started to be formed around the cells. By the end of this stage (at the point of 24 h after inoculation) dense clots were clearly visible in the ECM (Fig. 3**F**), which, in our opinion, should be considered as crystal nuclei that subsequently transform in the most thermodynamically stable form, calcite. It is interesting that the cells in this environment formed filaments joined with each other in long twisted chains and created a network, the nodes of which served as sites for the further active crystal growth. In 2015, van Gestel et al (31) carried out an elegant study, where they demonstrated intercellular interactions and division of labor during the migration of *B. subtilis* cells throughout biofilm formation. The researchers showed the enhancement of the expression of matrix-forming genes in cells at the edges of the colonies and organization of the cells into so-called van Gogh bundles consisting of filamentous loops. We have observed a similar picture for planktonic cells (Fig. 3): the cells united into long twisted chains, folded in stacks and, what was the most remarkable, an extracellular matrix has appeared around them. In their work, van Gestel et al suggested that the appearance of bending filaments was a response to external conditions. In our work, this was evidenced by the proteins required for matrix formation expression increase, in particular the aldehyde/alcohol dehydrogenase accumulated in experimental cells (Fig. 5**A** and **B**). Expression of the aldehyde/alcohol dehydrogenase *B. licheniformis* gene ortholog, which protein product catalyzes the reaction of acetyl CoA into ethanol necessary for matrix formation, has been reported to maximize during the induction of matrix formation in bacteria of other species (32).

The next conclusion we made was that the matrix not only served as a glue for binding cells to each other and as a site of the crystal growth, but also as a response to stress caused by conditions in the reaction medium (high pH, elevated concentrations of urea and calcium ions). It was evidenced by the accumulation of protease of the Clp family during the growth of *B. licheniformis* in the presence of urea and CaCl_2_ (Fig. 5**A** and **C**). We showed that urea acted not only as a source of carbonate ions generated by urease as described earlier for urease-driven bacterial induction of mineralization (33–35), but also as a denaturing agent causing matrix formation (Fig. 4**B**). On the one hand, for some bacteria, for example, *Staphylococcus aureus* (36) and *B. amyloliquefaciens* (37), *B. cereus* (32) and *B. subtilis* (38), representatives of Clp family ATPases were reported to be required for biofilm (and ECM) formation. On the other hand, the expression of Clp genes, which are highly conserved among bacteria, is a typical response to stress (for example, heat, chemical or osmotic shock) leading to the presence of proteins with a misfolding of the polypeptide chain (39). Besides urea, an increase in the production of the stress-associated Clp family proteins can be explained by the effect of used elevated concentration of calcium ions (22 mM), which was close to or exceeded the value typical for the induction of stress reactions (40). This idea was supported by atomic force and light microscopy images of *B. licheniformis* cells incubated in the urea-free Ca^2+^-containing medium resulted in induction of the ECM formation (Fig. 4**C**). Noteworthy, at the concentrations used of the present work, both CaCl_2_ and urea, as well as their mixture induced the formation of extracellular matrix during the first 24 hours of incubation (Fig. 4**B-D**).

We also demonstrated the presence of electron-dense inclusions inside bacteria under our experimental conditions (Fig. 6**B**). This observation remarkably agrees with similar data of other authors (29). In particular, by TEM analysis of *B. licheniformis* DB1-9 the group of Han with coauthors revealed the dark and denser spots inside the cells and within the cell EPS (29). This may indicate the calcium carbonate nucleation inside the bacterial cell. Moreover, the fact that the protein belonging to the porin family (ompA ortholog) was identified (in a mixture with glyceraldehyde-3-phosphate dehydrogenase), led us to another conclusion. Accumulation of this protein we reliably observed in experimental cells between 15 and 24 h after inoculation (Fig. 5**A** and **D**). Taken together data on the experimentally observed increase in the concentration of this protein and its potential in the formation of bacterial extracellular vesicles (41) could lead to a hypothesis about a possible transport of crystal precursors outside of the cell by vesicles that were described earlier for magnetotactic bacteria (42).

In conclusion, our results support the hypothesis (29) that CaCO_3_ nucleation occurs inside *B. licheniformis* cells and further crystal growth and polymorphic transformations occur in the extracellular matrix. We demonstrated that the spatial arrangement of the cells is important for the active crystal growth and is dependent on stress-induced environmental factors. The extracellular matrix forms a favorable microenvironment for crystal growth providing a high concentration of ions necessary for CaCO_3_ crystal aggregation, fixation, and stabilization.

## 4. MATERIAL AND METHODS

### 4.1 Bacterium growth conditions and sample preparations

The strain *Bacillus licheniformis* DSMZ 8782 used in the study was kept at −20°C in LB medium (peptone 10 g/L, yeast extract 5 g/L and NaCl 10 g/L) containing 20% (v/v) glycerol. Prior to each experiment, an overnight culture of *B. licheniformis* was prepared from frozen glycerol stock by culturing in LB at 37°C with an orbital shaker operated at 110 rpm. After the growth, cells were separated aseptically by centrifugation at 2000 g for 10 minutes and re-suspended in 2 mL of 0.9% NaCl to prepare the inoculum.

Routinely, to evaluate mineralizing ability of the strain *B. licheniformis* DSMZ 8782 during the growth in a liquid medium, the inoculum of the microorganism was transferred into the flasks with 100 mL of the standard B4 liquid medium containing: yeast extract, 2 g/L; glucose, 10 g/L; urea, 2.5 g/L; CaCl_2_, 2.5 g/L. Pre-sterilized solutions of urea and CaCl_2_ were added separately to the medium after autoclaving. In each case, bacterium was cultivated at 37°C with shaking at 110 rpm with periodic removing of aliquots. Withdrawn aliquots were cooled to the room temperature 25°C followed by measurements of pH, optical density at 600 nm and urease activity.

To analyze CaCO_3_ particles, precipitates were filtered on a vacuum filter with a pore size of 0.2 μm, dried at 37°C in a thermostat and analyzed by powder X-ray diffraction, optical stereoscopic and scanning electron microscopes and IR Fourier spectrometer (see below).

To assess the influence of the medium components on cell behavior during CaCO_3_ mineralization, 4 variants of standard B4 medium were used:

1. B4 without urea and CaCl_2_ (B4-C, control medium);
2. B4 with the addition of urea (2.5 g/L) and without CaCl_2_ (B4-U);
3. B4 with the addition of CaCl_2_ (2.5 g/L) and without urea (B4-Ca);
4. B4 with urea and CaCl_2_ (B4-UCa).

Preliminary prepared inoculum was transferred into the flasks with different media and cultivated for 48 hours at 37°C with shaking. Two-mL aliquots were withdrawn each 3 hours, cells were centrifuged and re-suspended in 1 mL of sterilized distilled water. Bacterial surface was studied using an NT-MDT Solver Bio scanning probe microscope in a semi-contact mode, using an NSG01 probe and by MSP-1 optical stereoscopic microscope.

### 4.2. Urease assay

Urease activity was determined according to the method described in (43) by evaluation of conductivity of a culture medium. To assess the urease activity, the aliquot was cooled to 25°C and the conductivity was measured using a conductometer Gravity: Analog Electrical Conductivity Sensor/Meter (K=10). The formation of ionic particles of non-ionic substrates led to an increase in the total conductivity of the solution and the rate, at which the conductivity increased, was proportional to the concentration of the active urease presented in the reaction mixture. The conductivity of the medium with growing *E. coli* DH5α strain was taken as a negative control. Molar concentration of degraded urea was calculated using the equation (1) with the coefficient taken from (43):

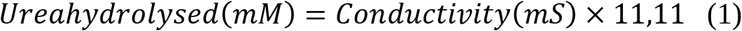

One unit of the urease activity corresponded to the enzyme amount capable of catalyzing the conversion of 1 μM urea per minute under the standard assay conditions (pH 5.5, 37° C, 20 min). The data points are presented as the means of at least three independent experiments, and the errors were calculated for each data point using Excel Solver add-in (Microsoft, Redmond WA). Reported below time-dependent curves for pH- and urease activity assays during the bacterial growth were generated with ORIGIN 8.0 software (OriginLab, Northampton, MA, USA).

### 4.3. CaCO_3_ particle analysis

Powder X-ray diffraction (XRD) analysis of the samples was performed on a Rigaku Miniflex 600 diffractometer (Bragg–Brentano geometry) with Ni-filtered CuKα (λ=1.5418Å) radiation and a LYNXEYE detector. Diffraction patterns were recorded in the 10–70° 2θ range, with a step of 0.02° and collection time of 0.3 s/step. Rietveld analysis of the patterns was carried out with a help of FullProf Suite (44). Structure models were obtained from Crystallography Open Database (45).

The microstructure of the samples was investigated using a Carl Zeiss NVision 40 high resolution scanning electron microscope at 1 kV acceleration voltage.

The IR spectra of samples were recorded on an ALPHA IR Fourier spectrometer (Brucker) in the disturbed total internal reflection mode in the range of 400–4000 cm^−1^ with a resolution of 1.5 cm^−1^.

### 4.4. Extracellular matrix isolation

To isolate fractions containing extracellular matrix (ECM), 2-mL aliquots were withdrawn from the 100-mL flasks containing medium B4-UCa or control B4-C (without urea and CaCl_2_) at 15 hours, 18 hours, 21 hours, and 24 hours during the growth of *B. licheniformis* at 37°C with shaking. Each sample was centrifuged at 8000 g for 10 minutes. Fractions containing supernatant (S1) were frozen at −20°C for further analysis; the pellets were re-dissolved in 2 mL of PBS and centrifuged for 10 minutes at 8000 g. The cell fraction containing extracellular matrix was frozen at −20°C for further analysis (fraction C). The number of cells was estimated by the suspension absorption at 600 nm using spectrophotometer Hitachi U-3310 and normalized. Cells were destroyed by ultrasound treatment followed by electrophoretic separation of proteins.

### 4.5. Extracellular matrix analysis

Samples of PBS-washed bacteria pellet (fractions C) were mixed with Laemmli’s buffer containing β-mercaptoethanol (46), denatured at 99°C for 5 minutes, and loaded into the wells of a polyacrylic gradient gel (8-16%) using a Kaleidoscope kit (BioRad) as a molecular weight marker. Fragments of the Coomassie-stained gel were excised and prepared for mass-spectrometric analysis as described in (47). Briefly, the gel fragments were washed from the dye twice with 30 mM NH_4_HCO_3_, 40% acetonitrile, dehydrated with 100% acetonitrile, and treated with trypsin (Promega) (20 μg/mL in 50 mM NH_4_HCO_3_) at 37°C for 5 hours. Tryptic peptides were mixed with a 2,5-DHB matrix (Bruker), applied to a target, and mass spectra were obtained on a MALDI-TOF/TOF mass spectrometer UltrafleXtreme (Bruker) in the positive ion mode. For each spectrum, 5000 laser pulses were summed up. Protein identification was performed using MASCOT (www.matrixscience.com) using the NCBI database (www.ncbi.nlm.nih.gov). The error was limited to 20 ppm. Methionine oxidation and deamidation were indicated as variable modifications. Identification was considered reliable (p <0.05) if the score value exceeded the threshold value.

### 4.6. TEM analysis

Bacteria-containing suspensions were centrifuged at 5000 g for 10 min and fixed with 2.5% PBS-buffered glutaraldehyde solution (Sigma Inc.) for 1 h. The glutaraldehyde-fixed bacterial cells were twice washed by PBS and post-fixed in 1% OsO_4_ solution for 1 h. Then, the cells were dehydrated in series of ethanol solutions of gradually increasing concentration, infiltrated with acetone and embedded in the Epon epoxy resine. The ultrathin sections (50-70 nm) of the bacteria under study were obtained using ultramicrotome Leica UC7 (Germany). The sections were collected on the 200 mesh copper grids and contrasted with uranyl-acetate and lead citrate. TEM analysis of the sections was performed using the electron microscope JEOL JEM 1011 (Japan), equipped with a high-resolution digital camera Morada (Olympus, Japan).

## ACKNOWLEDGMENTS

The authors acknowledge financial support from the Genome Research Center development program “Kurchatov Genome Center – PNPI” (agreement No. 075-15-2019-1663). This work was performed using the equipment of the Shared Research Center FSRC “Crystallography and Photonics” RAS and was supported by the Russian Ministry of Education and Science (project RFMEFI62119X0035). The SEM and IR-spectroscopy measurements were performed using shared experimental facilities supported by IGIC RAS state assignment.

